# The Effect of Celecoxib and muMab911 on Strain Adaptive Bone Remodeling and Fracture Repair in Female Mice: Implications for Rapidly Progressive Osteoarthritis

**DOI:** 10.1101/2025.03.24.645018

**Authors:** Nicholas Ruggiero, Alexandra Ciuciu, Ashkan Sedigh, Ibtesam Rajpar, David Shelton, Patrice Belanger, Kathryn Gropp, John A. Collins, Theresa A. Freeman, Ryan E. Tomlinson

## Abstract

Debilitating pain is the primary clinical feature of osteoarthritis (OA) that drives the enormous healthcare costs. Osteoarthritis-related pain is often treated with non-steroidal anti-inflammatory drugs (NSAIDs), which effectively relieve pain and inflammation by inhibition of prostaglandin synthesis. Antibodies directed against nerve growth factor (NGF) were tested some time ago as an alternative potential analgesic for musculoskeletal pain, including osteoarthritis-related pain. Unfortunately, clinical development of these drugs was put on hold due to adverse outcomes – primarily rapidly progressive osteoarthritis. Both prostaglandin synthesis and NGF have been implicated as critical mediators of strain adaptive bone remodeling, which may play a role in rapid osteoarthritis progression. Therefore, this study was designed to investigate the effects of celecoxib, an NSAID, and muMab911, an anti-NGF antibody, as well as the combination therapy on strain adaptive bone remodeling, bone mass and geometry, and bone healing in a murine model. Adult female C57BL/6J mice received celecoxib through drinking water, up to 3 IP injections of muMab911, or both treatments over a period of two weeks. As expected, all treatments were effective for relieving injury-associated pain. Consistent with previous studies, we found that celecoxib alone and in combination with muMab911 impaired periosteal load-induced bone formation induced by axial forelimb compression. Furthermore, both treatments had minimal effects on osteoblast and osteocyte populations, bone structural and material properties, and cortical and trabecular bone mass. Mice subjected to damaging axial forelimb compression did not have impaired healing or callus morphology due to treatment. Nonetheless, both drugs decreased NGF expression during fracture healing. In total, our results indicate that these medications both suppress NGF expression and load-induced bone formation which may support rapid OA disease progression by diminishing strain adaptive bone remodeling in subchondral bone of OA patients.

## INTRODUCTION

Osteoarthritis (OA) is a painful musculoskeletal condition involving the progressive loss of joint articular cartilage that affects nearly 600 million people worldwide [1]. Although age is the most significant risk factor for OA [2], the most clinically significant symptom of OA disease initiation and progression is joint pain [3]. Osteoarthritis-related chronic pain can range from intermittent discomfort to debilitating pain that severely impacts mobility and quality of life. As a result, there has been a widespread effort to determine the efficacy of existing FDA-approved medications against osteoarthritis-related pain as well as to develop novel therapeutic strategies that address this large clinical need.

Non-steroidal anti-inflammatory drugs (NSAIDs) effectively relieve joint pain and inflammation [4, 5]. NSAIDs act to prevent the synthesis of prostaglandins through the blockade of at least one of the cyclooxygenase (COX) enzyme isoforms, COX1 and COX2 [5]. Whereas the constitutively active COX1 generates prostaglandins that are required to maintain physiological functions, COX2 produces prostaglandins that drive pain, fever, and inflammation [6]. As a result, COX2-specific inhibitors, such as celecoxib, were introduced to minimize gastrointestinal bleeding and platelet abnormalities associated with non-specific NSAIDs [7, 8]. However, concern regarding serious adverse cardiovascular events have led to caution when prescribing COX2-specific inhibitors [9] as well as controversial follow-up clinical trials [10]. Moreover, NSAIDs often provide inadequate pain relief for patients with end-stage osteoarthritis [11].

To address this unmet clinical need, research has focused on nerve growth factor (NGF) as a key mediator of bone and joint pain and a potential therapeutic target [12–15]. Early studies showed anti-NGF therapy effectively relieved OA, fracture, and cancer pain [12, 16], though it had no beneficial effect on cartilage or synovium [17]. Indeed, treatment with anti-NGF antibodies was found to significantly diminish pain from new-onset and established osteoarthritis [16]. To date, three monoclonal antibodies have been developed for use in humans. Tanezumab was used for the first large randomized, placebo-controlled trial for relief of osteoarthritis pain by anti-NGF therapy [15]. In that study, patients on tanezumab experienced a large reduction in knee pain while walking (up to 62%), with only minor common adverse events, such as headache. However, unexpected adverse events from Phase II and III trials resulted in a clinical hold [17]. In particular, tanezumab-treated patients had more joint safety events, specifically rapidly progressive osteoarthritis (RPOA) [18], and required more total joint replacements than placebo-treated patients [19]. Lower doses still increased joint safety events, including RPOA which was linked to NSAID use and may result from neuropathic arthropathy [18–21]. To date, the mechanism by which anti-NGF antibodies, particularly in conjunction with NSAID usage, can cause RPOA remains poorly understood [20].

We and others have shown that both COX2 and NGF signaling are critically required by the skeleton to respond normally to mechanical load [21, 22]. Furthermore, inadequate strain adaptive bone remodeling plays an important role in osteoarthritis disease progression. In particular, early-stage OA involves decreased subchondral bone plate thickness and subchondral trabecular bone mass due to increased bone remodeling rate, whereas late-stage OA is associated with increased subchondral bone plate thickness, microdamage, and sclerotic trabecular bone due to diminished bone resorption [23]. Thus, strain adaptive bone remodeling appears to be the best explanation for the subchondral trabecular bone sclerosis observed in late-stage OA [24]. In total, we reason that inhibition of strain adaptive bone remodeling with NSAIDs and/or NGF antibodies would favor OA progression, particularly in patients with late-stage OA.

This study was designed to determine if these medications have a synergistic effect to diminish strain adaptive bone remodeling, affect bone material properties, or bone repair. In particular, we examine the effects of celecoxib, anti-NGF antibodies, and the combination treatment following damaging and non-damaging axial forelimb compression. We hypothesized that these pain relief medications would diminish strain adaptive bone remodeling and decrease skeletal repair, predisposing users towards OA disease progression and increasing the risk of RPOA.

## METHODS

### Mice

All experiments involving mice were approved by the Institutional Animal Care and Use Committee (IUCAC) of Thomas Jefferson University (Protocol #01919). Female C57BL/6J mice were purchased from the Jackson Laboratory (Stock #000664) at 15 weeks old and acclimated in our animal facility for two to three weeks until 17-18 weeks old. Mice were fed standard rodent chow (LabDiet 5001) and housed in groups of up to five.

### Medications

Mice were randomly separated into four treatment groups with 10 mice per group: vehicle control, celecoxib only, muMab911 only, and combination treatment of celecoxib and muMab911. The vehicle control group received ddH_2_O. The celecoxib only and combination treatment group received celecoxib through drinking water by dissolving celecoxib (40 mg/kg) in ddH_2_O to a final concentration of 250 mg/L with drinking water replenished every 2-3 days. Assuming an intake of approximately 4 mL of water per day per mouse, the average daily dose of celecoxib administered was 40 mg/kg. This dose represents approximately half the recommended maximum daily human dose (6.6 mg/kg), adapted to mice using allometric scaling provided by the FDA Center for Drug Evaluation and Research. Celecoxib drinking water was administered 24 hours before the start of experiments on day 0. Mice receiving muMab911 were administered IP injections of the anti-NGF antibody at 10 mg/kg on days 0, 6, and 12 (non-damaging loading) or days 1 and 7 (damaging loading) [25]. This dose corresponds with the human equivalent Tanezumab dose of 833 µg/kg, which is many multiples higher than the therapeutic range of Tanezumab (10-200 µg/kg) [15]. In all loading experiments, mice given combination treatment were dosed exactly as mice receiving celecoxib or muMab911 individually, but the treatments were administered concurrently.

### Mechanical Loading

Mice were subjected to mechanical loading using standard axial forelimb compression to induce lamellar bone formation (LBF) or woven bone formation (WBF) at the ulnar mid-diaphysis, as in previous work [21, 22, 26–28]. Before loading experiments, mice received 0.12 mg/kg of buprenorphine by IP injection and were anesthetized with inhaled isoflurane (2-3%) throughout the loading procedure. Deep anesthesia was confirmed by hindpaw pinch and then the mouse’s right forelimb was secured in a custom fixture for loading using a standard material testing system (TA Instruments Electroforce 3200 Series III).

For LBF experiments, mice received a 0.3 N compressive preload followed by 100 cycles of a 2 Hz rest-inserted sinusoidal waveform with a peak force of 3.0 N three times per week for a total of six bouts of LBF loading over two weeks. This peak force has been previously shown to cause an average peak compressive strain of approximately 3000 µɛ at the ulnar mid-diaphysis of C57BL/6J mice that induces significant LBF without causing damage to the bone [29, 30]. After loading, mice were allowed unrestricted cage activity and tissues were harvested on day 14.

For WBF experiments, loading parameters were chosen as a function of ultimate force and total displacement to fatigue fracture [28, 31]. To determine ultimate force, monotonic compression to failure by displacement ramp (0.1 mm/s) was used on representative mice outside of the treatment groups. To determine displacement to fatigue fracture, a sinusoidal waveform was applied to the representative mouse forelimb at 2 Hz and 75% of the ultimate force, which was determined to be 3.6 N for this group of mice, until complete fatigue fracture. The total displacement was calculated and 66% of the total displacement was used to generate a stress fracture at the ulnar mid-diaphysis, which was 0.84 mm for this group of mice. Thus, for WBF experiments, mice received a 0.3 N compressive preload followed by a cyclic sinusoidal waveform with a peak force of 3.6 N at a frequency of 2 Hz until a total displacement of 0.84 mm relative to the 10^th^ cycle. This protocol has been previously shown to consistently generate a non-displaced stress fracture at 1 mm distal to the ulnar midpoint, resulting in robust woven bone formation [28]. After loading, mice were allowed unrestricted cage activity and tissues were harvested on day 8, which was 7 days after injury on day 1.

### MicroCT

For LBF experiments, loaded and non-loaded forelimbs were harvested, dissected free of soft tissue, fixed overnight in neutral buffered formalin (NBF), washed in PBS, and stored in 70% ethanol until microCT scanning. Forelimbs were scanned separately using a Bruker Skyscan 1275 microCT system with a 1 mm aluminum filter, using 55 kV and 181 µA with a 73 ms exposure time and a 10 µm isometric voxel size. Transverse plane scan slices were obtained by placing the forelimb parallel to the z-axis of the scanner. Reconstruction and analysis were performed using NRecon and CTAn (Bruker), respectively. Outcomes measured focused on bone mass, density, and geometry. For WBF experiments, loaded (fractured) and non-loaded (intact control) forelimbs were harvested 7 days after injury, dissected free of soft tissue, and stored in PBS-soaked gauze at −20°C until microCT scanning. Forelimbs were scanned separately, reconstructed, and analyzed as the forelimbs from LBF experiments. Outcomes measured included woven bone formation, density, extent, and callus size.

In both LBF and WBF experiments, non-loaded femurs were harvested, dissected free of soft tissue, stored in PBF-soaked gauze at −20°C until microCT scanning. Femurs were scanned using 80 kV and 125 µA with a 47 ms exposure time and a 13 isometric voxel size. Again, transverse plan scan slices were obtained by placing the femur parallel to the z-axis of the scanner. Reconstruction and analysis were performed as previously described. Trabecular bone geometry was analyzed across a 1 mm region of the metaphysis that was 0.5 mm below the growth plate and cortical bone geometry was analyzed across a 1 mm region at the mid-diaphysis.

### Three-Point Bending

The structural and material properties of non-loaded femurs were analyzed using standard three-point bending. Femurs that were previously scanned using microCT and stored in PBS-soaked gauze at −20°C were thawed. Once at room temperature, the femurs were oriented on a standard fixture of a material testing system (TA Instruments Electroforce 3200 Series III) with femoral condyles facing down and a bending span of 7.29 mm. Then, a 0.3N preload was applied to fix the femur in place followed by a monotonic displacement ramp of 0.1 mm/s until failure while force and displacement values were recorded digitally. The outputted force-displacement curves were converted using microCT-based geometry and a custom GNU Octave script to stress-strain curves.

### Forelimb Mechanical Testing

For WBF experiments, microCT scanned loaded (fractured) and non-loaded (intact control) forelimbs were hydrated in PBS at 4°C overnight, brought to room temperature, then placed into the original fixtures of the material test system (TA Instruments Electroforce 3200 Series III) that were used for fatigue loading. Forelimbs were subjected to monotonic compression to failure at 0.1 mm/sec, a protocol previously shown to effectively measure structural properties of woven bone in a healing fracture [26]. The force-displacement data was used to calculate the strength, stiffness, and total energy to failure of the forelimbs using a custom MATLAB script and normalized to the contralateral limb.

### Finite Element Analysis

For LBF experiments, loaded and non-loaded forelimbs were scanned using microCT and 3D solid models of each bone were generated. Using the FEBio software package [32], finite element analysis of these forelimbs was conducted. A voxel-based finite element mesh of tetrahedral elements was created in PreView using Tetgen. Homogenous and isotropic material properties were assigned to the forelimb with an elastic modulus of 20 GPa and a Poisson’s ratio of 0.3 [33]. The model was constrained at the olecranon but permitted to displace in the Z dimension (distal/proximal axis) at the carpus to simulate *in vivo* axial forelimb compression. Circumferential strains at the midshaft were compared between treatment groups.

### Dynamic histomorphometry

To measure bone formation rates after LBF loading, calcein (10 mg/kg) and alizarin red (30 mg/kg) were IP injected into mice 2 and 7 days before harvesting tissues. Forelimbs were fixed overnight in NBF, dehydrated in graded ethanol, and embedded in polymethylmethacrylate (PMMA). Forelimbs were sectioned at the mid-diaphysis of the ulna with a low-speed diamond wafering saw (Isomet 1000) at a thickness of 100 µm, mounted on glass slides with Eukitt, and polished to a thickness of about 30 µm for imaging and analysis by dynamic histomorphometry. Images were taken using a Zeiss LSM 800 confocal microscope and analyzed for periosteal (Ps) and endosteal (Es) mineralizing surface (MS/BS), mineral apposition rate (MAR), and bone formation rate (BFR/BS) as defined by the ASBMR Committee for Histomorphometry Nomenclature [34]. Relative (r) parameters are calculated as the difference between loaded and non-loaded limbs.

### Sanderson’s Rapid Bone Stain

Thick PMMA-embedded sections used for dynamic histomorphometry in LBF experiments were subsequently stained using Sanderson’s Rapid Bone Stain (Dorn & Hart) for static histomorphometry measurements. These stained sections were analyzed for osteoblast number per bone surface and osteocyte number per bone area.

### Picrosirius Red Staining and Second Harmonic Generation Imaging

To analyze fibrillar collagen in LBF experiments, tibias from all treatment groups were harvested, dissected free of soft tissue, fixed overnight in NBF, decalcified in 14% EDTA for 7 days, and embedded in paraffin. Tibias were sectioned at 4 µm thickness and stained using picrosirius red for 1 hour according to the manufacturer’s instructions (Polysciences, #24901). Sections were then mounted with Eukitt onto microscope slides and imaged with a circularly polarized light microscope (Nikon Eclipse LV100POL). Using this method, the birefringence color changes from shorter to longer wavelengths (from green to red) as collagen fiber thickness increases [35]. Best-fit color thresholds were determined using individual pixel selection, with each pixel categorized as a red (thick), yellow (medium), or green (thin) fiber using NIS-Elements AR 4.5.00 (Nikon). Afterwards, sections were imaged with a two-photon microscope equipped with a 25x water immersion objective (Olympus FV1000MPE) at 860 nm with 5% laser power for second harmonic generation imaging. In both analyses, the entire tibial cross-section was quantified. Images were tiled and analyzed using FIJI [36].

### ELISA Assays

To analyze circulating markers of bone remodeling in LBF experiments, blood was harvested from the mice immediately after euthanasia via cardiac puncture to collect blood serum. The blood was allowed to coagulate for 30 minutes before being centrifuged at 5000g for 10 minutes to isolate serum. Serum was stored at −80°C until used and ELISA kits against receptor activator of nuclear factor kappa-B ligand (RANKL), osteoprotegerin (OPG), procollagen type 1 intact N-terminal peptide (P1NP), and serum cross-linked C-telopeptide of type I collagen (CTX), along with a standard ELISA kit to measure erythropoietin levels, were used to quantify circulating levels of these proteins (RANKL: R&D Systems #MTR00; OPG: R&D Systems #MOP00; P1NP: Novus Biologicals #NBP276466; CTX: Novus Biologicals #NBP3-11802; EPO: R&D Systems #MEP00B). Serum was diluted 2-fold for RANKL and 5-fold for OPG and diluted samples were analyzed in triplicate and compared to a linear standard curve. A four-parameter logistic standard curve was created using Curve Expert (Hyams Development).

### Protein Quantification Wes

For WBF experiments, two additional groups of female C57BL/6J mice were subjected to stress fracture to analyze protein concentrations of nerve growth factor (NGF) in loaded (fractured) and non-loaded (intact control) forelimbs 1 or 3 days after fracture. After harvesting the forelimbs, the middle third of the ulna was placed into TRIzol (Life Technologies), pulverized in liquid nitrogen using a freezer/mill (SPEX 6575), and processed following the manufacturer’s instructions. Protein was isolated from pulverized bone powder by dialyzing (Spectra/Por 7 RC - MWCO 2000) the phenol-ethanol supernatant obtained using the manufacturer’s instructions (Trizol) against 0.3% SDS for 24 hours. Protein concentration was assessing using BCA assay (Thermo), then samples were prepared by dilution in 0.1x Sample Buffer to obtain a concentration of 1.0 mg/mL, then mixing with 5x Fluorescent Master Mix (1:4 ratio). Next, the diluted samples were denatured at 95 °C for 5 min. Primary antibody against NGF (Abcam ab6199) was diluted in Antibody Diluent 2 (ProteinSimple), then the biotinylated ladder (5 uL), diluted samples (3 uL), primary antibodies (10 uL), secondary antibodies (10 uL), luminol-peroxide (15 uL), and wash buffer (500 uL) were loaded into the WES plate. The entire plate was centrifuged (1000g for 5 minutes) before starting the assay. The capillary electrophoresis results were analyzed and quantified using Compass for Simple Western software.

### Forelimb Asymmetry

Following WBF loading, behavioral tests were done on days 0, 3, 4, 5, 6 and 7 to determine stress fracture-related pain levels and analgesic effects of the administered treatments. Forelimb asymmetry tests were done by placing the mouse inside a clear, cylindrical tube with two mirrors positioned at angles to reflect the tube backside, and a camera recording the entire cylinder for five minutes. After filming, mice are removed and returned to their cage. Each incidence of vertical exploration where the mouse stood on its hindlegs and supported its body weight by placing its forepaws on the tube was scored to quantify forelimb use or aversion. A score of 1 was given if only the right (injured forelimb) forepaw was used, 0.5 if both forepaws were used, and 0 if only the left (uninjured forelimb) forepaw was used, as in previous work [37]. A minimum of 10 and a maximum of up to 42 independent vertical exploration incidents were recorded and compared to pre-injury baselines (D0).

### Von Frey Filament Testing

Additionally, mechanical hyperalgesia of the stress fracture limb was tested using Von Frey filament testing. Mice were acclimated for several hours in a plexiglass chamber with a metal mesh floor on at least 2 of the 5 days before Von Frey testing was performed. On testing day, mice were given 30 minutes of resting time in the plexiglass chamber, then testing began using the lowest calibrated Von Frey filament followed by higher forces. The paw plantar surface of the stress fracture limb was probed 5 times for 2 seconds with at least 5 seconds between probes and the paw withdrawal threshold was determined to be the force at which withdrawal occurred in at least 60% of the tests. After reaching the threshold, the next highest Von Frey filament was used to confirm that the threshold was reached.

### Study Design & Statistics

Preliminary data and data from previous studies were used to estimate the sample sizes for each treatment cohort using power analyses for the outcome with the highest variability, dynamic histomorphometry, to achieve 90% power at the 0.05 level of significance [21]. The calculated sample size was 9 mice per treatment group, so 10 mice were used to account for potential attrition. After data collection and unblinding during statistical analysis, outliers were identified using Grubbs’ test. In statistical tests that compared loaded to non-loaded forelimbs, the loaded forelimbs were normalized using non-loaded forelimb data before comparisons between treatment groups. For other analyses, raw data was compared to data from vehicle treated controls. Statistical analysis was performed using Prism 10 (GraphPad Software Inc.). At each time point, a one-way ANOVA was performed across treatment groups and all analyses were performed without matching using Dunnett’s multiple comparison correction with an adjusted p-value of less than 0.05 considered significant.

## RESULTS

### Celecoxib, muMab911, and combination treatment all decrease load-induced bone formation

Load-induced bone formation in mice subjected to six bouts of non-damaging axial forelimb compression over a period of two weeks was assessed using standard dynamic histomorphometry (**Fig. 1A-D**). Representative images from each treatment group illustrates that loading significantly increased periosteal bone formation in loaded limbs (**S. Fig 1**) as compared to non-loaded limbs (*not shown*). Specifically, treatment with celecoxib significantly decreased both periosteal MS/BS (−36%) and MAR (−47%), leading to a significant decrease in BFR/BS (−49%) in loaded limbs as compared to vehicle (**Fig. 1E-G**). In contrast, treatment with muMab911 alone did not significantly affect either periosteal MS/BS or MAR, but a strong trend (p = 0.051) was observed in BFR/BS (−27%) in loaded limbs as compared to vehicle (**Fig. 1E-G**). Finally, the combination treatment of celecoxib and muMab911 resulted in a similar effect as celecoxib alone, with significant decreases in both periosteal MS/BS (−55%) and MAR (−30%), resulting in a significant decrease in BFR/BS (−48%) in loaded limbs as compared to vehicle (**Fig. 1E-G**). No significant differences were observed in endosteal MS/BS, MAR, or BFR/BS with treatments (**Fig. 1H-J**). In total, these results illustrate that celecoxib and muMab911 both decrease periosteal load-induced bone formation to varying degrees, but the combination treatment does not result in additive effects.

**Figure 1.**
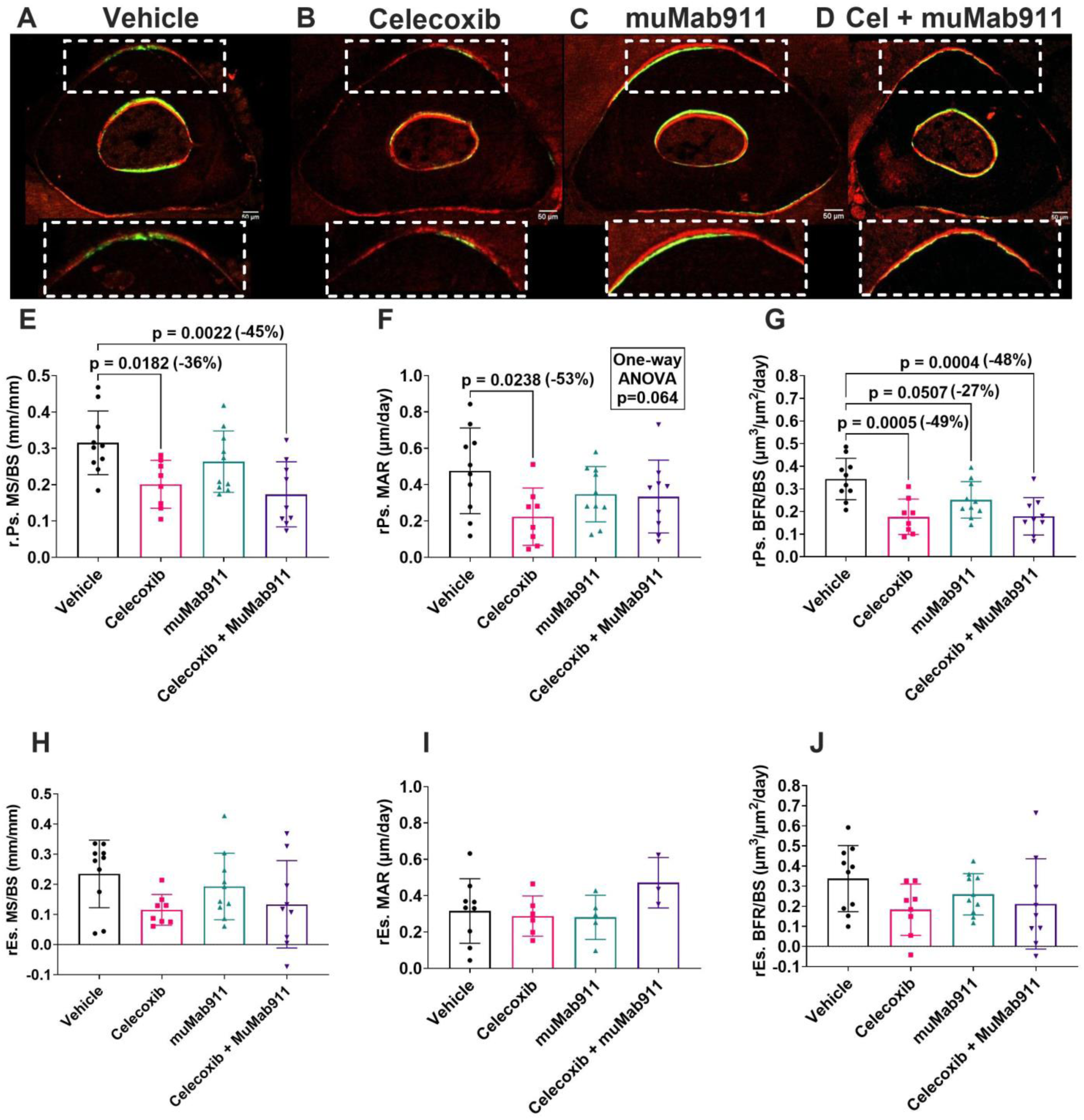
Celecoxib and celecoxib with muMab911 diminish load-induced bone formation on the periosteal surface as compared to vehicle controls. A-D) Load-induced bone formation was quantified by dynamic histomorphometry using mid-ulnar cross-sections with calcein and alizarin red bone formation labels. Relative measures were calculated as the loaded limb minus the non-loaded limb value. Outcomes include (E) relative periosteal mineralizing surface per bone surface (r.Ps.MS/BS), (F) relative periosteal mineral apposition rate (rPs.MAR), (G) relative periosteal bone formation rate per bone surface (r.Ps.BFR.BS), and (H-J) associated endosteal measures. Significance = p < 0.05 vs. vehicle, n=9-10 per group.

### Treatments do not change osteoblast and osteocyte populations in the ulna

Changes in the response of bone to mechanical forces could be caused by drug-induced changes in osteoblasts or osteocytes that are mechanoresponsive cells in bone [21, 38]. To address this potential treatment effect, the number of osteoblasts per bone perimeter and the number of osteocytes per bone area were quantified at the ulnar mid-diaphysis in SRBS-stained PMMA sections (**S. Fig 2**). However, no significant differences in osteoblast or osteocyte populations between treatment groups were observed (**Fig. 2**). These data indicate that the observed changes in strain adaptive bone remodeling are not due to a cytotoxic effect of the respective treatments on bone cells *in vivo*.

**Figure 2.**
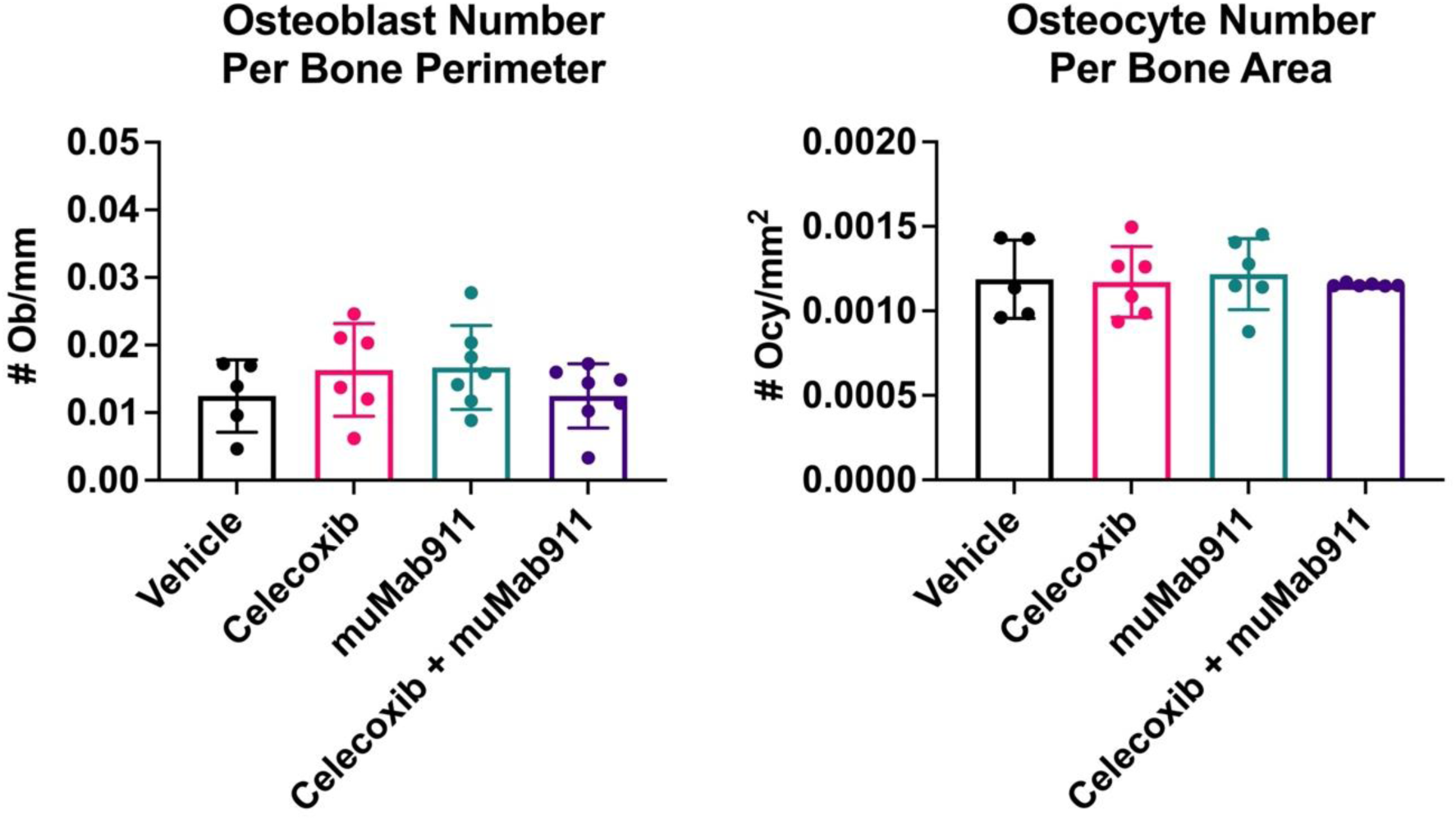
Treatments do not affect osteoblast and osteocyte populations in the ulna. Osteoblast number per bone perimeter and osteocyte number per bone area were measured on PMMA-embedded sections of the midshaft ulna. N = 5-7 per group.

### Treatments have minimal effects on forelimb strain distribution

Finite element analysis was used to examine if changes in strain adaptive bone remodeling caused by drug treatment resulted in altered strain distribution in the forelimb. In our analysis, we observed no significant differences in strain between treatment groups in either the non-loaded or loaded ulnas separately (**Fig. 3A,B, Fig. 3D,E**). When considering strain of the loaded (right) limb relative to the non-loaded (left) limb, we observed a small increase (+12%) in relative midshaft strain with celecoxib treatment (**Fig. 3C**) and a slight decrease (−15%) in relative max strain with combination treatment (**Fig. 3F**). In total, these results suggest that these treatments have minimal effects on forelimb strain distribution.

**Figure 3.**
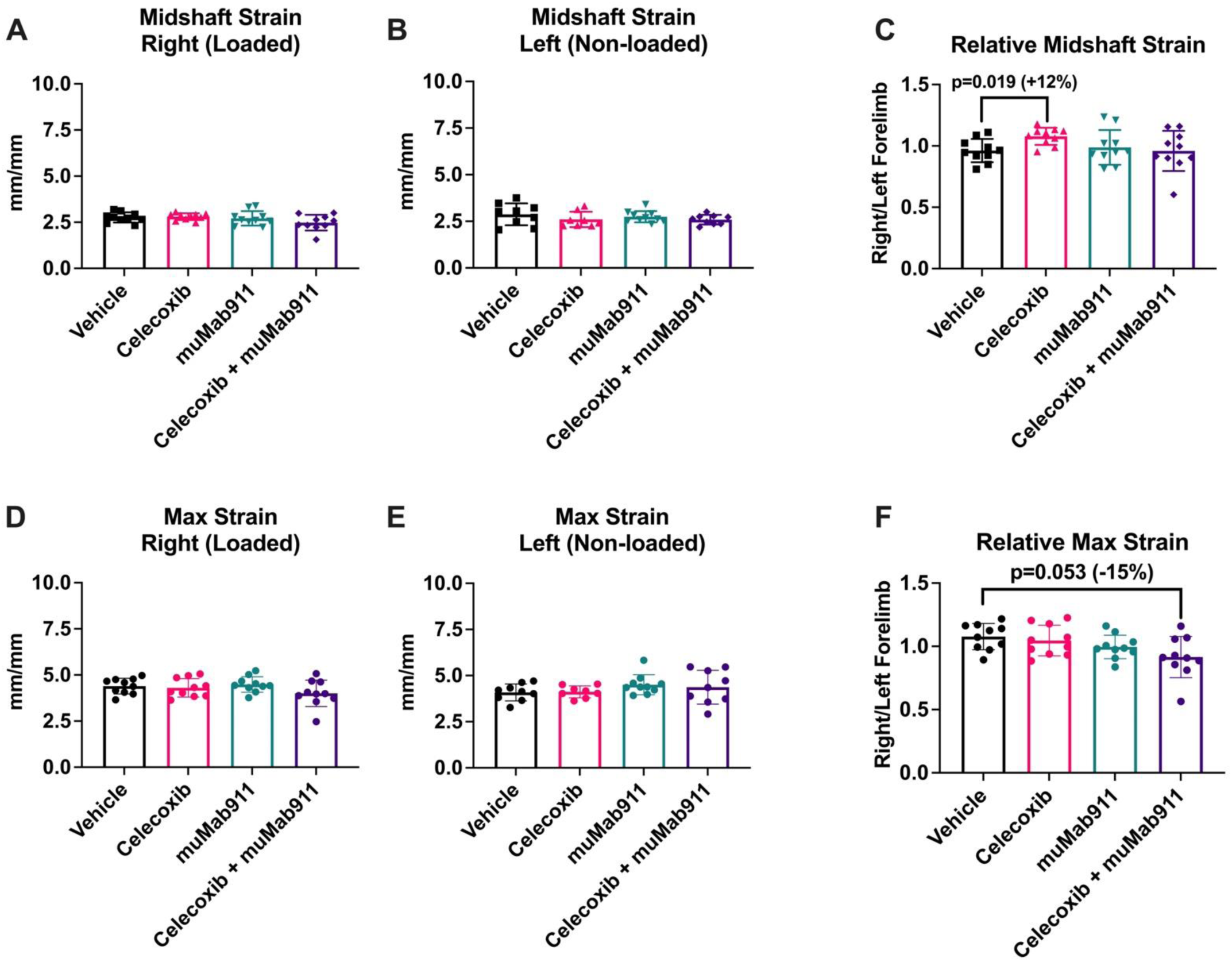
Celecoxib and celecoxib with muMab911 have significant, but small, changes on midshaft and max strain in the ulna when modeled with finite element analysis. (A-B) Right (loaded) and left (non-loaded) ulna midshaft strain quantifications by finite element analysis using FEBio with a mesh of tetrahedral elements created from microCT-based geometry using Tetgen. Each model was subjected to unit displacement in the Z axis with unrestricted movement at the carpus and restriction at the olecranon, then analyzed at the midshaft. (C) Relative ulna midshaft strains of right relative to left limbs. (D-E) Right (loaded) and left (non-loaded) ulna max strain quantifications. (F) Relative ulna max strains of right relative to left limbs. Significance= p<0.05 vs. vehicle, n=8-10 per group.

### Treatments have modest effects on trabecular and cortical bone

Next, we utilized microCT of non-loaded femurs to analyze the effects of treatment on bone mass and geometry (**Fig. 4**). Trabecular bone geometry parameters were not strongly affected by treatment (**Fig. 4E-H**); the only significant difference was a minor increase (+12%) in trabecular number in combination treatment bones as compared to vehicle (**Fig. 4G**). Surprisingly, we observed a small (−1%) but significant decrease in cortical tissue mineral density (TMD) in all treatment groups relative to vehicle (**Fig. 4I**). However, there were no differences in cortical bone geometry (**Fig. 4J-L**). In total, these results are consistent with previous studies illustrating that a short course of these medications minimally affect bone mass and geometry [22].

**Figure 4.**
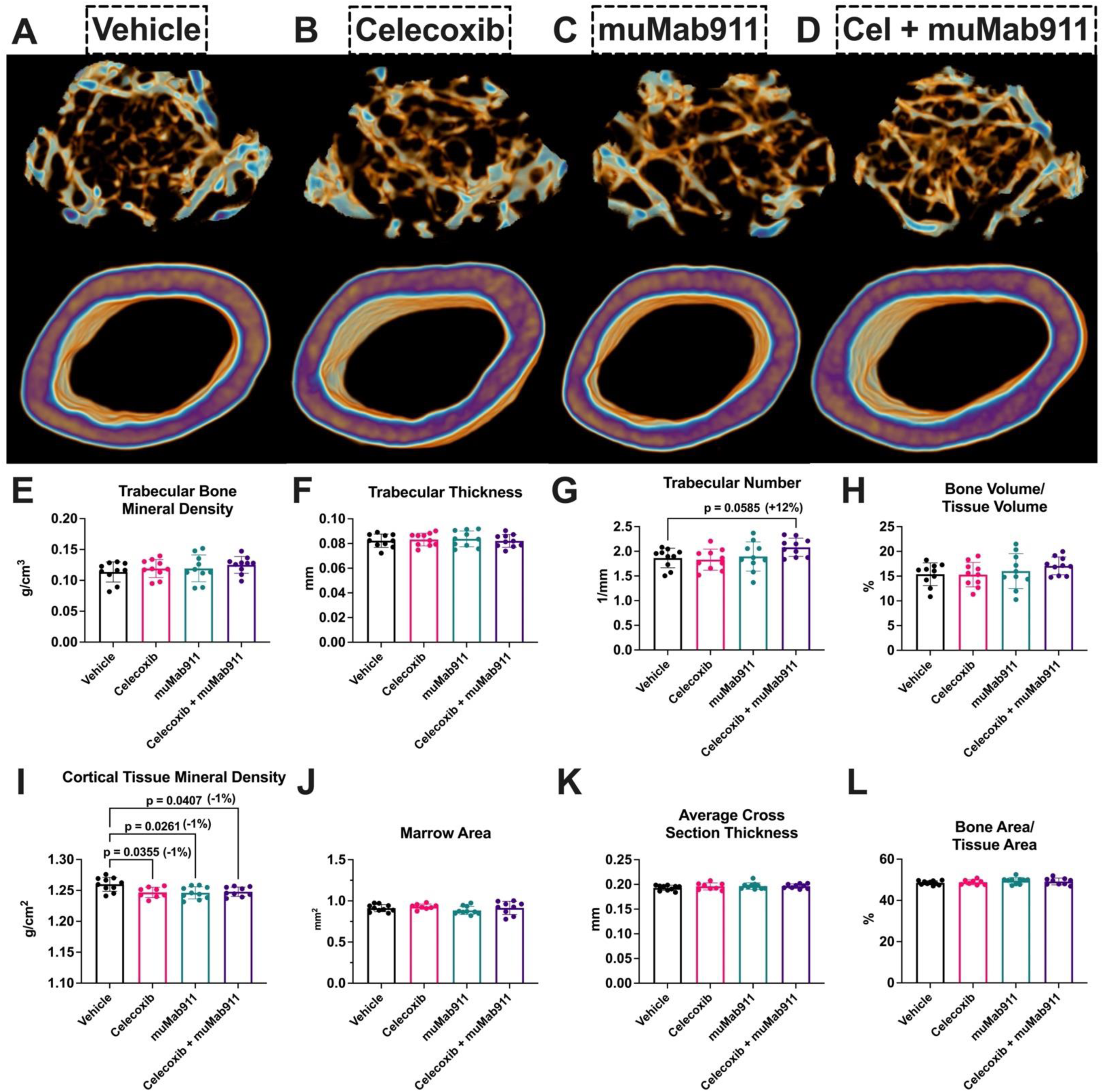
Celecoxib has minor affects on trabecular geometry, but all treatment groups have comparable cortical geometry to vehicle mice. (A-D) Representative reconstructions of trabecular and cortical regions of interest for vehicle, celecoxib, muMab911, and combination treatment groups as measured by microCT. Quantifications of trabecular bone mineral density (E), trabecular thickness (F), trabecular number (G), trabecular bone volume/tissue volume (H), cortical tissue mineral density (I), marrow area (J), average cortical cross section thickness (K), and cortical bone area/tissue area (L). Significance= p<0.05 vs. vehicle, n=8-10 per group.

### Treatments do not affect structural and material properties of the femur

In a previous study, we observed a significant loss in bone toughness in female C57BL/6J mice receiving the NSAID naproxen through a mechanism related to collagen fibril thickness [22]. First, we determined if these medications similarly affect bone structural and material properties by subjecting non-loaded femurs to standard three-point bending. Here, no significant differences due to treatment were detected in any parameters, including ultimate bending moment, bending rigidity, ultimate stress, Young’s modulus, energy to failure, or toughness (**Fig. 5**). Consistent with these findings, we observed no significant differences due to treatment in collagen fiber thickness as quantified by picrosirius red stained sections (**S. Fig 3**) or signal from second harmonic generation (SHG) imaging (**Fig. 6**). Therefore, these data demonstrate that celecoxib, muMab911, and the combination do not affect bone toughness or collagen organization in bone.

**Figure 5.**
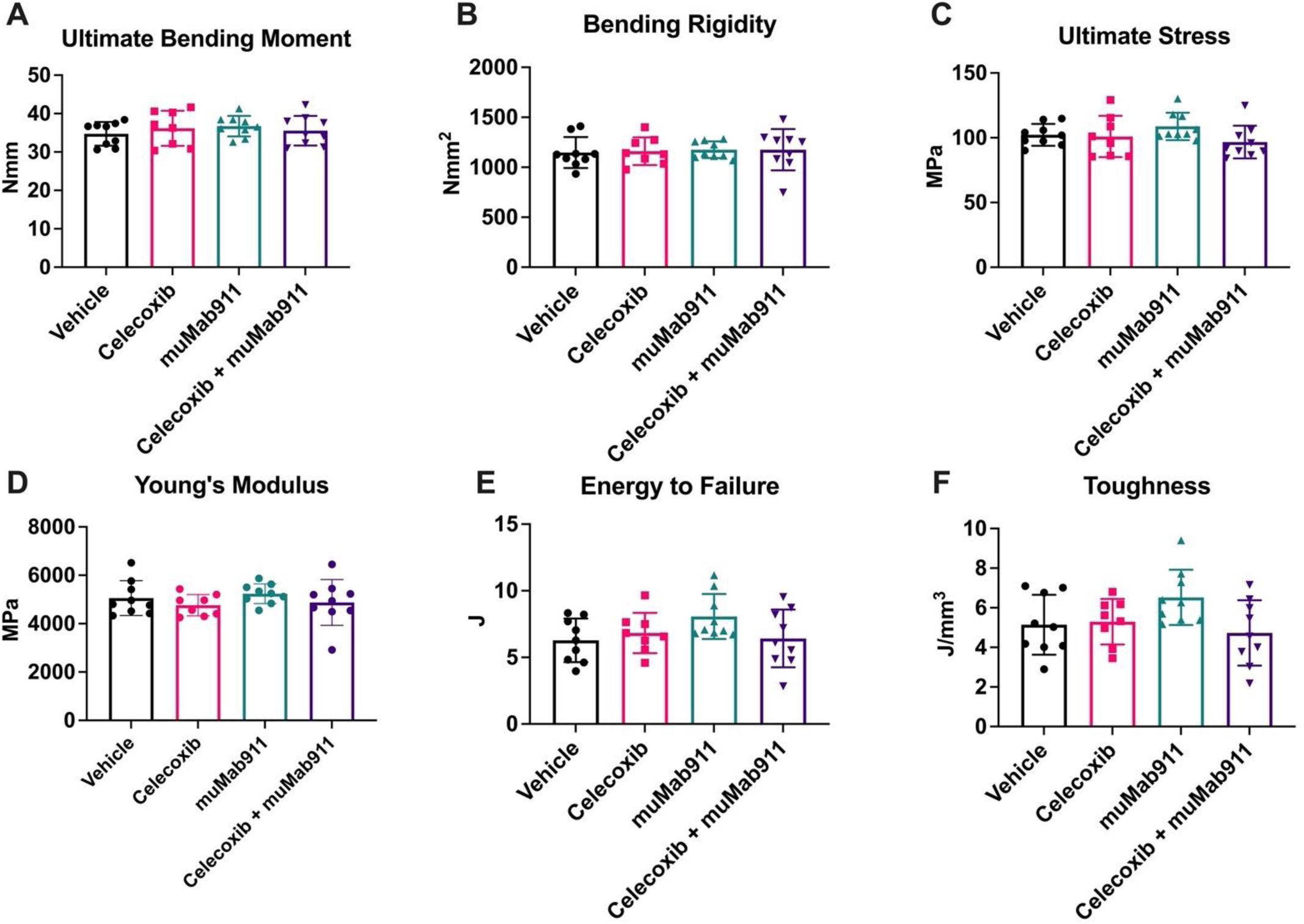
No treatment group affects bone structural or material properties as compared to vehicle controls. Standard three-point bending of non-loaded femurs was done to determine (A) ultimate bending moment, (B) bending rigidity, (C) ultimate stress, (D) young’s modulus, (E) energy to failure, and (F) toughness. Significance= p<0.05 vs. vehicle, n=6-8 per group.

**Figure 6.**
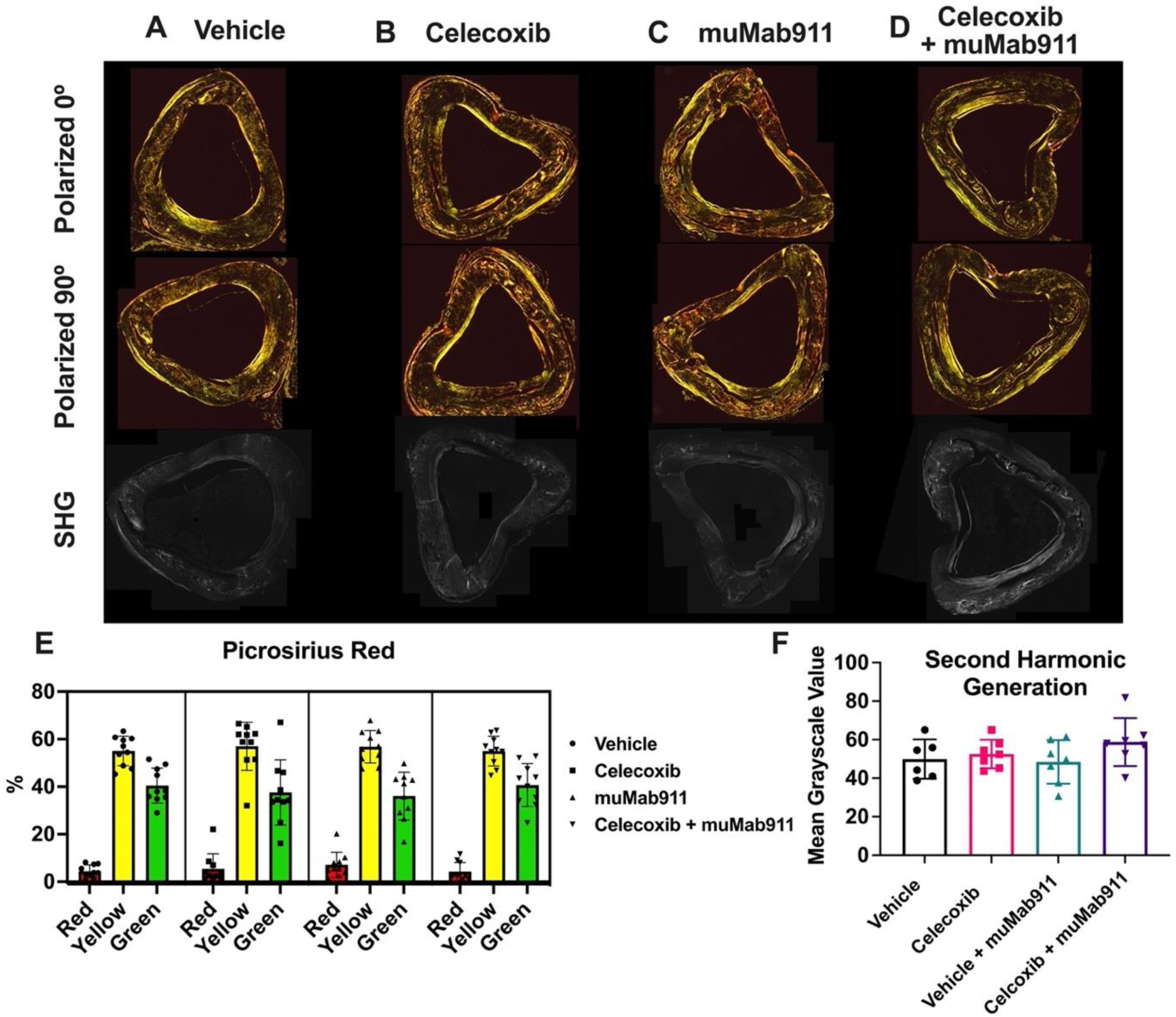
No changes to tibial collagen organization are observed with celecoxib, muMab911, or combination treatments. (A-D) Represnetative images of picrosirius red stained sections visualized with polarized light and second harmonic generation (SHG) imaging of vehicle, celecoxib, muMab911, and combination groups. (E) Quantification of the percentage of red (thick), yellow (medium), and green (thin) collagen fibers in tibial cross-sections. (F) Quantification of the average second harmonic generation signal in the entire tibial cross-section. Significance= p<0.05 vs. vehicle n=6-10 per group.

### Effect of treatments on serum bone biomarkers

Next, we assessed potential systemic effects of treatment using serum that was collected from mice at end term following LBF loading. Using ELISA assays, no significant differences between treatments were observed in the bone formation marker P1NP (**Fig. 7A**). Surprisingly, there was a significant increase in the bone resorption marker CTX with muMab911 treatment (+40%; p=0.0427), though this increase was not present in the combination treatment group (**Fig. 7B**). No significant differences in levels of OPG, RANKL, or the ratio of RANKL to OPG were observed between treatments (**Fig. 7C-E**). Finally, mice receiving muMab911 alone had a trend (p=0.085) of increased erythropoietin in their serum as compared to vehicle that was not significant in post-hoc testing (**Fig. 7F**). In all markers measured, the combination treatment did not have any additive effects on protein concentration in serum. In addition to these serum bone biomarkers, celecoxib concentration was also quantified in serum following one or two weeks of treatment (**S. Fig. 4**). Here, celecoxib concentration was below the lower limit of detection at one week in all treatment groups and was only significantly elevated from baseline in the combination treatment group at two weeks.

**Figure 7.**
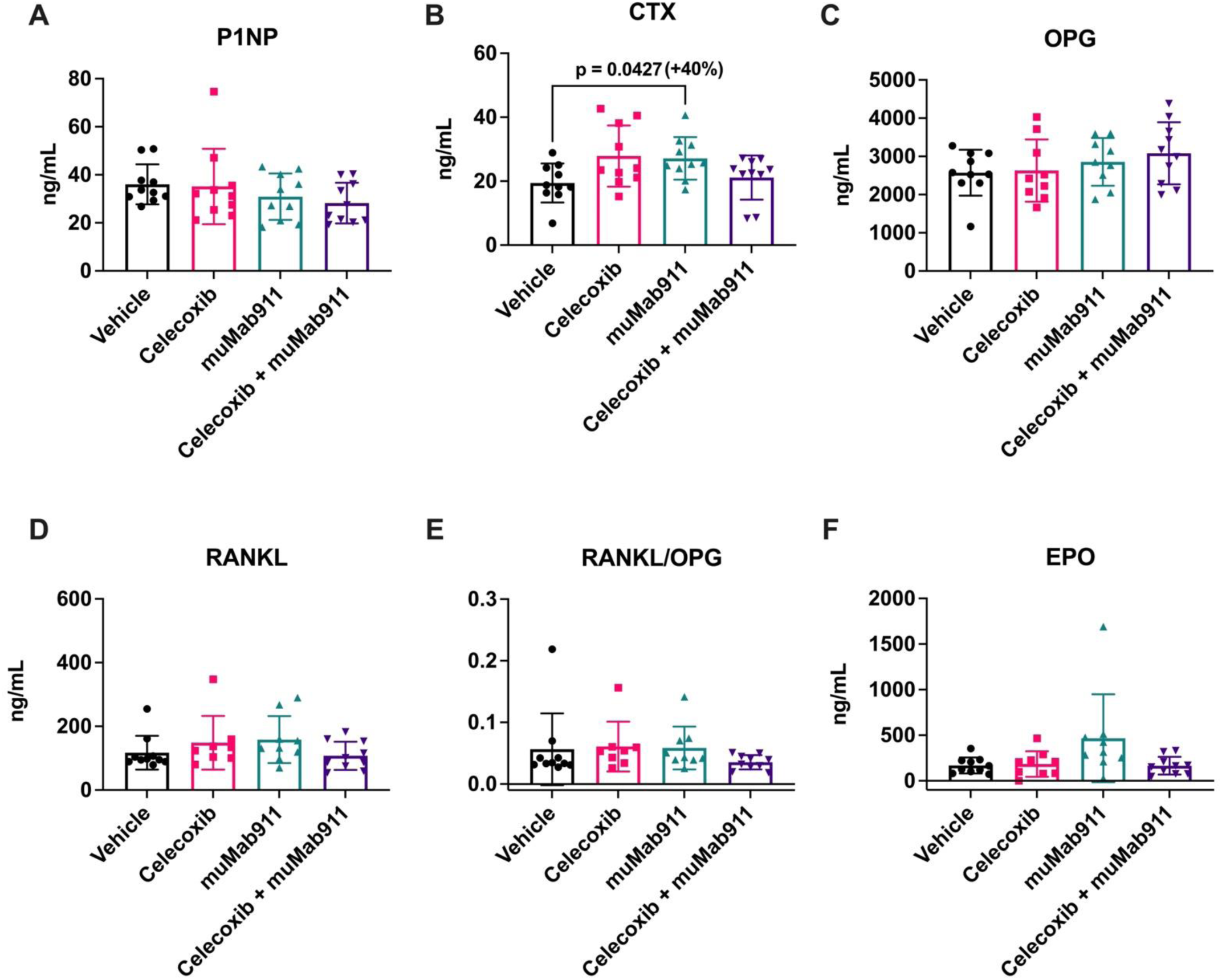
muMab911 significantly increases circulating CTX and shows a trending increase in erythropoietin. Serum levels as measured by ELISA of (A) P1NP, (B) CTX, (C) OPG, (D) RANKL, (E) RANKL/OPG, and (F) EPO. Significance= p<0.05 vs. vehicle, n=8-10 per group.

### Treatments do not affect fatigue fracture formation or callus formation after injury

First, we observed that 24 hours of pre-treatment with celecoxib, muMab911, or combination treatment did not significantly change the number of cycles required to induce fatigue fracture, defined as displacement of 0.84 mm relative to the 10^th^ cycle (**Fig. 8A**). The effects of treatment on fatigue fracture healing were assessed using microCT seven days post-injury. Here, we found no significant change due to treatment in any quantified measure of the callus (**Fig. 8B-G**). Regarding woven bone volume, there was an apparent decrease with combination treatment as compared to vehicle that was not observed with either celecoxib or muMab911 alone, but this effect was not statistically significant (p=0.25). Additionally, the similar woven bone and crack extent demonstrate that there were similar levels of damage during fracture induction between different treatment groups. Overall, these data indicate that these doses of celecoxib and muMab911 used do not impede fatigue fracture healing seven days after injury, and that the combination treatment was not additively detrimental to healing in this pre-clinical model of fatigue injury.

**Figure 8.**
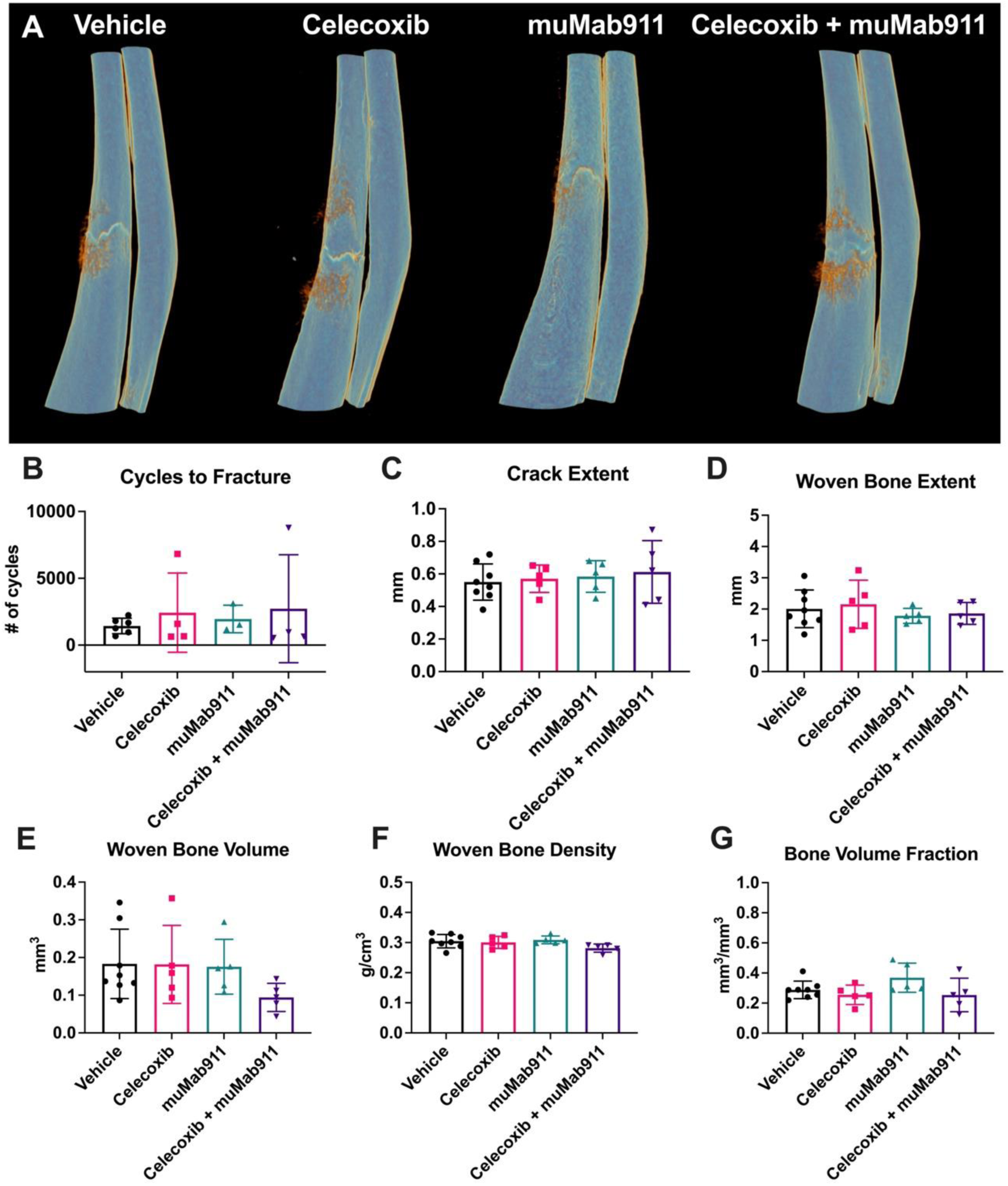
Celecoxib, muMab911, or combination treatment do not affect the number of cycles to fatigue fracture formation or fracture healing seven days post-injury. (A) Representative reconstructions of fatigue fracture calluses seven days post-injury in vehicle, celecoxib, muMab911, and combination treatment groups. (B) The number of cycles required during WBF experiments for the forelimb to reach the goal displacement of 0.84 mm. After seven days of healing, the woven bone callus of mice within each treatment group were analyzed by microCT for (C) crack extent, (D) woven bone extent, (E) woven bone volume, (F) woven bone density, and (G) bone volume fraction of tissue volume. Significance= p<0.05 vs. vehicle, n=5-8 per group.

### Treatments relieved fatigue fracture-related pain and NGF expression in early fracture healing

First, the analgesic efficacy of each drug was assessed using forelimb asymmetry tests on days 3 to 7 after stress fracture. Here, mice receiving any treatment tended to use their injured (ipsilateral) forelimb more often than vehicle treated mice, with a strong trend of forelimb usage normalized to baseline (p=0.060). Of all treatment groups, muMab911 showed the highest injured forelimb usage relative to baseline as compared to vehicle, particularly on 6 and 7 days after injury (**Fig. 9A**). Von Frey filament testing was also performed to determine the forepaw withdrawal threshold for each forelimb. The results from day 0 (before fracture) and day 3-7 (after fracture) in the non-loaded (intact) forelimb relative to loaded (fractured) forelimb illustrate that vehicle treated mice had a lower withdrawal threshold than all treatment groups at day 7; thus, all treatments effectively relieved pain (**Fig. 9B**). Finally, the protein levels of nerve growth factor (NGF) were measured in the stress fracture injured bone of a separate cohort of female C57BL/6J mice 1 or 3 days after stress fracture injury. Results from 1 day after injury showed that vehicle mice had significantly higher levels of NGF at the injury site than mice receiving any treatment (**Fig. 9C**). Surprisingly, celecoxib alone and the combination treatment decreased NGF levels more than the anti-NGF antibody, muMab911. In contrast, quantification of NGF 3 days after injury revealed no significant differences between groups (**Fig. 9D**).

**Figure 9.**
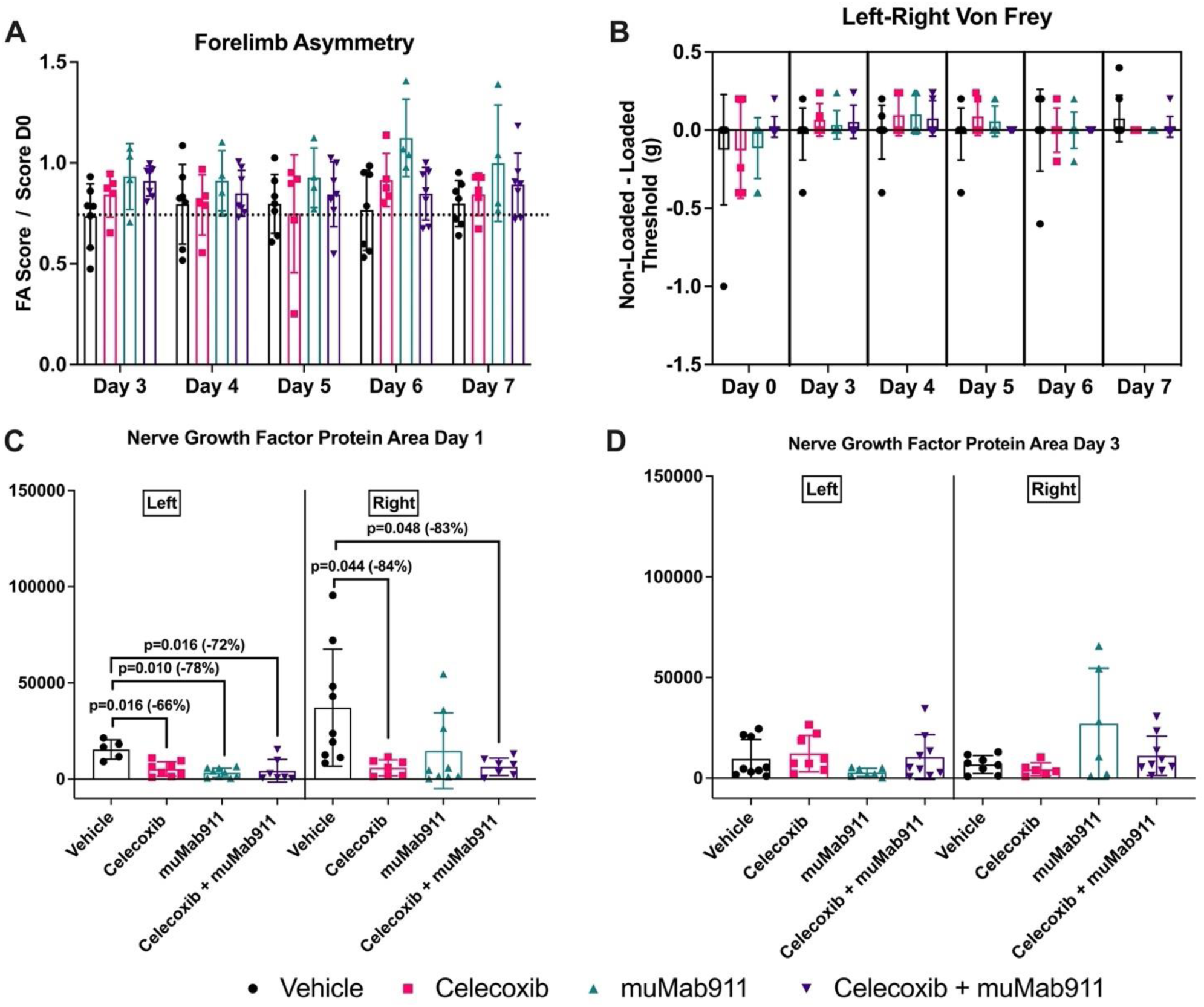
Effects of celecoxib, muMab911, or combination treatment on forelimb usage and NGF protein levels after fatigue fracture. Baseline (D0) forelimb usage scores per group measured before injury were used to normalize scores after injury for each group. (A) Forelimb asymmetry quantifications relative to baseline levels per treatment group, n=5-7 per group. (B) Von Frey withdrawal thresholds of left (non-loaded) limbs relative to right (fractured) forelimbs, n=5-9 per group. (C-D) NGF protein levels in injured ulna midshaft 1 and 3 days after injury as measured by ProteinSimple Wes and divided between left (non-loaded) and right (fractured) forelimbs, n=5-9 per group. Significance= p<0.05 vs. vehicle.

### Effect of treatments on stress fracture repair

The effect of each treatment on stress fracture repair was assessed by performing monotonic axial compression of injured and contralateral (control) forelimbs until failure. Here, there were no significant differences due to treatment in the control forelimbs, indicating that the treatment alone for eight days did not affect the mechanical performance of the bones (**Fig. 10A-C**). Similarly, there were no significant differences in these values due to treatment in the fractured forelimbs (**Fig. 10D-F**). However, when fracture limbs were analyzed relative to their own intact contralateral forelimb, we observed significant differences in ultimate load, where celecoxib treated bones displayed a significant increase (+79%) as compared to vehicle (**Fig. 10G**); a similar effect that was not statistically significant was observed in the combination group. Although there were no significant differences between treatment groups in stiffness (**Fig. 10H**), ultimate energy was significantly different between treatment groups with both celecoxib (+121%) and combination (+118%) significantly increased as compared to vehicle. (**Fig. 10I**).

**Figure 10.**
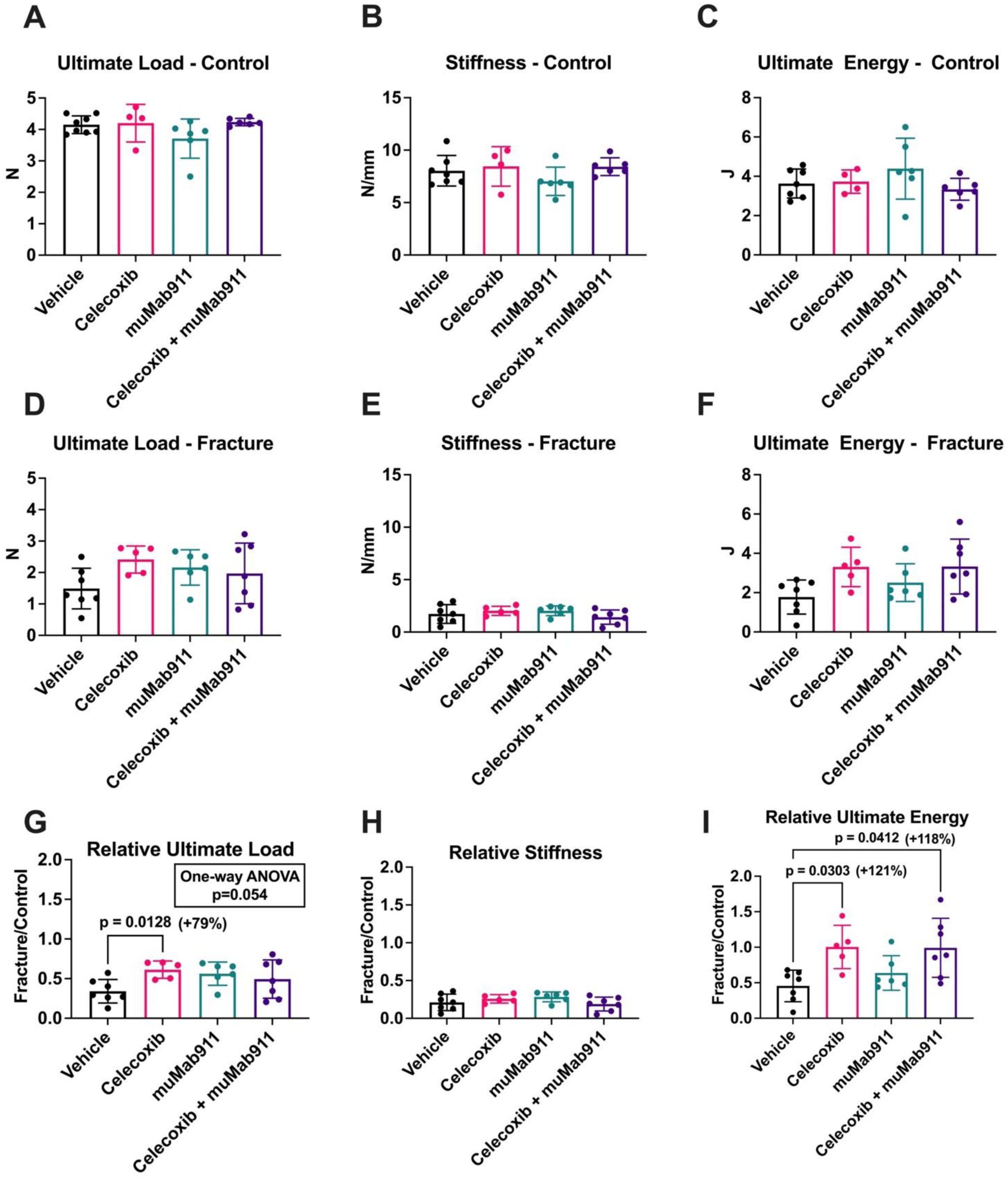
Celecoxib and combination treatment significantly increase the mechanical performance of healed limbs relative to intact limbs. (A-C) Ultimate load, stiffness, and ultimate energy measurements of left (intact, non-loaded) forelimbs subjected to monotonic compression until failure in the same set up as used for fatigue fracture induction. (D-F) Ultimate load, stiffness, and ultimate energy measurements of right (woven bone) forelimbs seven days after injury. (G-I) Ultimate load, stiffness, and ultimate energy measurements of right forelimbs normalized to left. Significance= p<0.05 vs. vehicle, n=5-7 per group.

## DISCUSSION

Rapidly progressive osteoarthritis (RPOA) induced by anti-NGF therapy has been well-described, but its etiology is poorly understood and there is no distinguishing pathophysiology. Furthermore, clinical trials of the anti-NGF antibody tanezumab demonstrate a dose-dependent increase in joint safety events when compared to continuing a stable dose of NSAIDs in patients with hip or knee osteoarthritis [39]. Therefore, this study was designed to uncover additive deleterious effects of anti-NGF and NSAIDs therapy, particularly in the response of the skeleton to mechanical force that may underpin structural changes in late-stage osteoarthritis [24]. Here, we found that administration of the NSAID celecoxib or anti-NGF antibody muMab911 both diminished load-induced bone formation in mice (**Fig. 1**), consistent with previous studies [21, 22], but did not display an additive effect. Taken together with our observation in this study that celecoxib is more potent than muMab911 to prevent load-induced bone formation, these results suggest that impaired strain adaptive bone remodeling is unlikely to explain the increased incidence of RPOA due to concurrent NSAID usage with anti-NGF therapy.

Our previous report demonstrated that two weeks of treatment with the NSAIDs naproxen and aspirin did not affect bone mass or geometry in mice [22]. Surprisingly, we did observe that naproxen significantly diminished bone toughness through effects on collagen fibril thickness. In this study, we observed limited effects of celecoxib and muMab911 on bone mass, geometry, or strength (**Fig. 4-5**) as well as no significant differences in collagen fibril thickness as indexed by picrosirius red staining (**Fig. 6**). To our knowledge, this is consistent with the literature; no previous reports have indicated effects of celecoxib or muMab911 on collagen fibril thickness. However, we did observe a small but significant effect of diminished cortical bone tissue mineral density (TMD) in all three treatment groups. Additional studies are required to determine the mechanism by which these medications uniformly affect this important parameter related to bone strength.

Furthermore, we did not observe impaired stress fracture healing associated with either celecoxib or muMab911 treatment in this study. We had hypothesized that celecoxib would negatively affect fracture repair, as the preponderance of studies have found that NSAIDs are deleterious for fracture repair [40], including several studies utilizing celecoxib in various animal models [41–43] and our own recent study examining the effects of naproxen on stress fracture healing in mice [22]. We speculate that this inconsistency with previous results may be due to poor delivery of celecoxib, potentially due to the strong hydrophobicity of celecoxib in water, based on our serum celecoxib concentration (**S. Fig 4**). Taken together with the efficacy of celecoxib against stress fracture-related pain, these results suggest that lower, intermittent dosages of celecoxib may effectively relieve pain and diminish strain adaptive bone remodeling without affecting fracture healing outcomes.

Finally, the requirement of NGF-TrkA signaling for fracture repair has previously been investigated. In one study, the loss of TrkA signaling using a chemical-genetic knockout mouse model significantly inhibited stress fracture repair in mice [44]. In contrast, blockade of NGF-TrkA signaling via neutralizing antibodies against NGF and TrkA reduced pain but did not influence healing of an osteotomy gap stabilized by an external fixator in mice [45]. Taken together with our findings, these results suggest that bone healing is governed by NGF-TrkA signaling but inhibition of this signaling during the healing cascade using neutralizing antibodies is limited. As a result, anti-NGF therapy may be appropriate for pain relief during fracture repair, although future studies should directly assess this outcome.

This study did have several limitations. In particularly, this study only used young adult, female C57BL/6J mice that were exposed to relatively short pain relief treatments. It is relevant to consider extending this work towards males and older mice, as this may be relevant from a translational standpoint. Furthermore, administration of celecoxib through drinking water may have suffered from a common drawback of this drug delivery method – inconsistent delivery of celecoxib in treated animals. Furthermore, our results suggest that potential interactions between celecoxib and muMab911 may prolong celecoxib in serum (**S. Fig. 4**), increasing the potential for adverse side effects with longer treatment. As a result, future studies may consider an alternative bolus dosing strategy. Finally, our finite element analysis did not incorporate the alterations observed in cortical tissue mineral density with regards to strain distribution.

## CONCLUSION

This study demonstrates that administration of celecoxib, muMab911, or combination treatment impairs load-induced bone formation compared to vehicle treatment but does not negatively impact bone structural and material properties, stress fracture formation or healing. Additionally, it was observed that many significant differences due to treatment occurred in celecoxib alone or combination treatments, suggesting that the inhibition of COX2 may drive any negative effects of NSAIDs used in combination with anti-NGF antibodies. Although significant differences in measures such as circulating markers of bone resorption were observed with celecoxib and muMab911 treatment, the effect sizes were generally small. This pre-clinical study of an NSAID and anti-NGF antibody in female mice indicates that these drugs in conjunction do not significantly increase the risk of injury or impede healing in the context of non-damaging and damaging forelimb loading. Additionally, results of celecoxib or muMab911 alone indicate that these drugs alone are not likely to suppress microdamage repair of subchondral bone in osteoarthritis patients, but both may impair load-induced bone formation in subchondral sclerotic bone that could work in conjunction at the minimal therapeutic doses to increase incidences of rapidly progressing osteoarthritis as previously observed clinically.

## ACKNOWLEDGEMENTS

This study was funded through a cooperative research agreement by Pfizer, Inc. DS, PB, and KG were Pfizer employees when this work was planned and conducted but are not currently.

## AUTHOR CONTRIBUTIONS

Conceptualization: RET. Investigation: NR, AC, AS, IR, RET. Analysis: NR, AC, RET. Supervision: DS, PB, KG, JC, TAF, RET. Writing – Original Draft: AC, JC, TAF, RET.

## SUPPLEMENTAL FIGURES

**Supplemental Figure 1.**
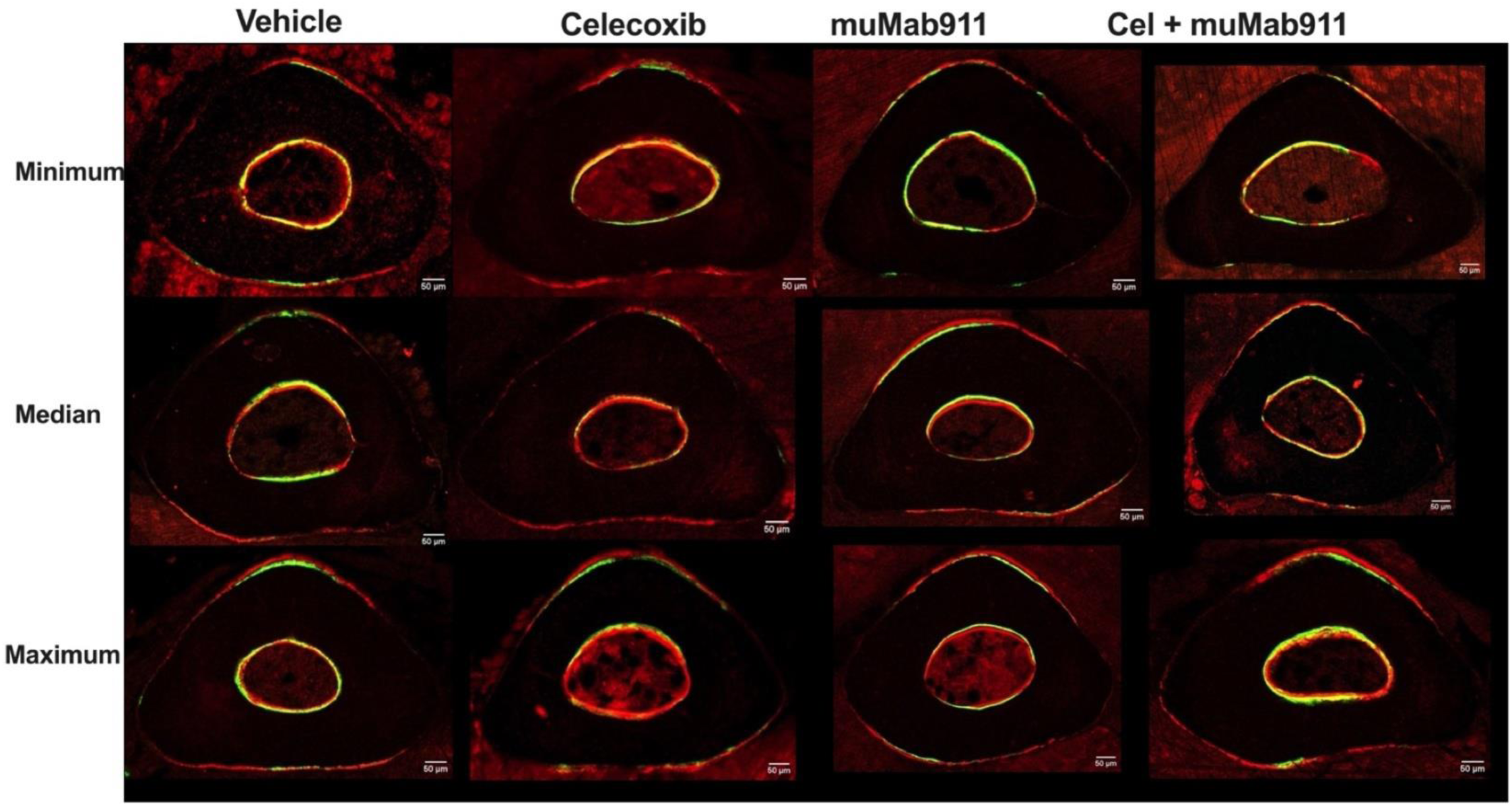
Representative images of minimum, median, and maximum sections from dynamic histomorphometry quantifications for each treatment group. Representative images were identified using individual periosteal bone formation rate/bone surface.

**Supplemental Figure 2.**
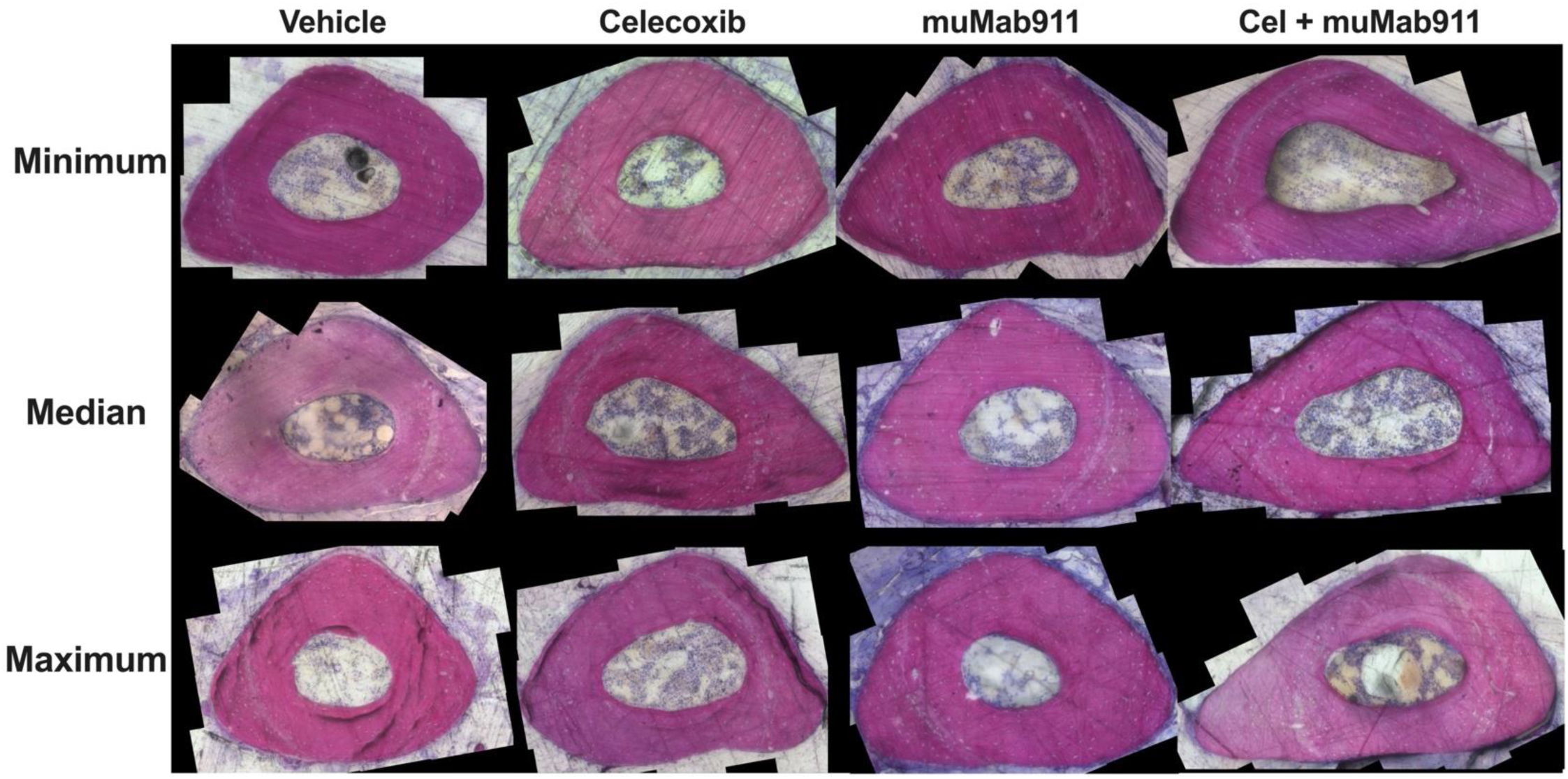
Representative images of minimum, median, and maximum sections from SRBS quantifications for each treatment group. Representative images were identified using individual osteocyte number.

**Supplemental Figure 3.**
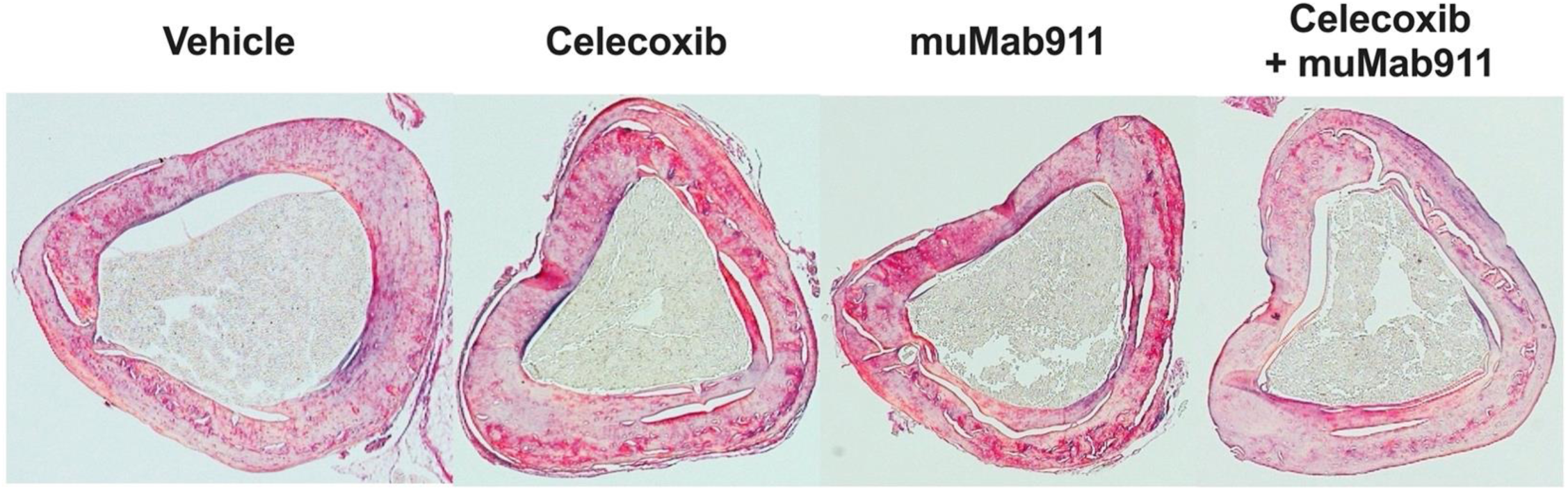
Representative brightfield images of picrosirius red-stained tibias. Representative images were identified using mean values for red, yellow, and green birefringence.

**Supplementary Figure 4.**
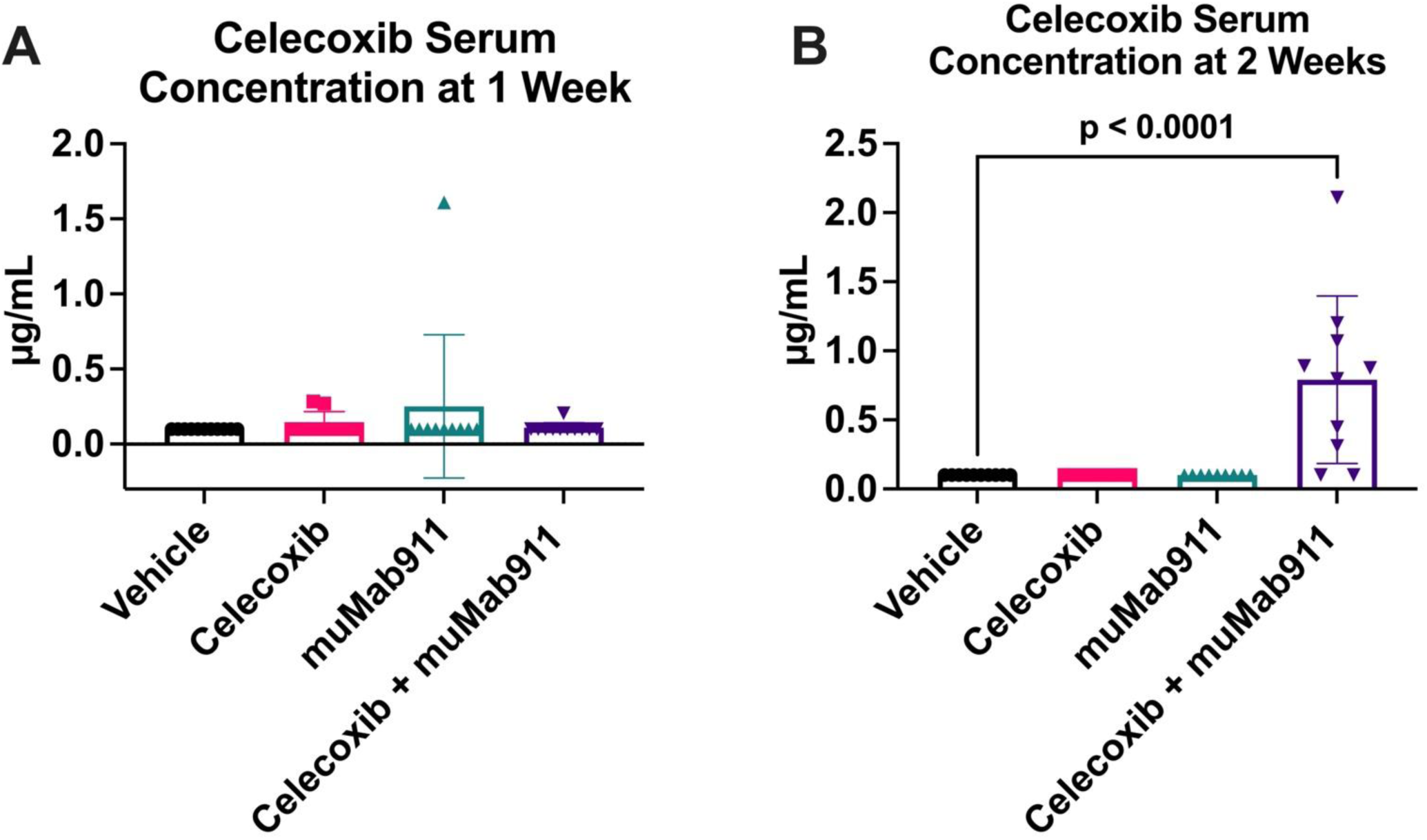
Celecoxib concentration in blood serum varied greatly between treatment groups and timepoints. (A) Quantification of celecoxib in blood serum after one week of treatment. (B) Quantification of celecoxib in serum after two weeks of treatment.

